# Differential D_1_ and D_2_ receptor internalization and recycling induced by amphetamine *in vivo*

**DOI:** 10.1101/2022.06.10.493955

**Authors:** Hanne D. Hansen, Martin Schain, Helen P. Deng, Joseph B. Mandeville, Bruce R. Rosen, Christin Y. Sander

**Affiliations:** Neurobiology Research Unit and Center for Integrated Molecular Brain Imaging, Copenhagen University Hospital, Rigshospitalet; Copenhagen, Denmark; Athinoula A. Martinos Center for Biomedical Imaging, Department of Radiology, Massachusetts General Hospital; Charlestown, MA, USA; Harvard Medical School; Boston, MA, USA

## Abstract

The dopamine system plays a significant role in drug reward and the pathogenesis of addiction. Psychostimulant drugs acutely increase dopamine levels, triggering receptor internalization. *In vitro* data suggest that dopamine D_1_ receptors (D_1_R) recycle, whereas D_2_ receptors (D_2_R) degrade in response to activation. Yet, receptor fates *in vivo* remain unclear. This study bridges *in vitro* mechanisms and *in vivo* measurements of stimulant-induced modulation of receptor states using longitudinal multi-modal imaging combined with neuropharmacology. We demonstrate how repeated amphetamine administration differentially modulates D_1_R vs. D_2_R signaling in nonhuman primates over 24 hours using simultaneous positron emission tomography and functional magnetic resonance imaging. In contrast to predominantly inhibitory D_2_R signaling due to an initial amphetamine challenge, excitatory D_1_R functional signaling prevails three hours later, while D_2_Rs stay internalized. These results demonstrate differential externalization mechanisms of the D_1_R and D_2_R *in vivo* and a shift in receptor subtype activation after a dopamine surge.

## INTRODUCTION

Substance use disorders are characterized as the progressive loss of control from initial and voluntary drug intake with reinforcing and hedonic effects to loss of control. This behavior becomes habitual and eventually compulsive. According to the World Drug Report 2021, an estimated 0.5% of the global population, or 27 million people, use amphetamine-type stimulants, with the highest prevalence in North America at 2.3% *(1)*. The non-medical use of stimulants has substantial medical, social, and economic consequences.

It is well-recognized that dopamine (DA) dysregulation accompanies addictive behavior. Data from *in vivo* animal and human studies reveal that, although stimulant drugs acutely increase DA levels in the striatum and reinforce their rewarding effects, reduced DA signaling is associated with behavioral features that facilitate the development and severity of addiction long-term [reviewed by Trifilieff et al. *(2)*]. Specifically, significant reductions in DA release, DA transporter availability, and dopamine D_2_ receptor (D_2_R) availability have been found in chronic stimulant users *(3)*. Despite much progress from *in vivo* receptor measurements, relatively little is known about the interplay of other DA receptor subtypes with D_2_Rs and their signaling mechanisms during repeated stimulant exposure in the living brain. Such mechanisms may play an important role in drug reward and the formation of addiction *(4, 5)*.

Amphetamine-type stimulants act on DA transporters and presynaptic vesicles to increase extracellular DA *(6, 7)*. This drug-induced increase in synaptic DA can trigger receptor internalization as one of the immediate responses to adapting to overwhelmingly high concentrations of DA *(8)*. Receptor internalization is considered an essential mechanism for discharging the bound agonist and making receptor sites available again on the surface of the cell membrane *(9)* – a homeostatic adaptive process at the receptor level to downregulate functionality during DA surges. More than half of the D_2_Rs can undergo internalization upon exposure to high concentrations of agonist *(10, 11)*. Furthermore, receptor internalization is mediated by β-arrestin2 *(12)*, and the genetic elimination of this protein in knock-out animals causes changes in the behavioral responses to most classes of drugs of abuse *(13). In vitro*, it has been found that both dopamine D_1_ receptors (D_1_R) and D_2_R rapidly internalize in response to DA release *(14, 15)*. However, the intracellular fate of dopaminergic receptor subtypes may be quite different: A study by Bartlett et al. found that the D_1_R quickly recycles back to the cell membrane, whereas the D_2_R is degraded *(16)*. The latter is in agreement with the observation that D_2_Rs, once internalized, can stay internalized for several hours or days after a single stimulant exposure *(17)*. The possibility that D_1_Rs may be available for binding by DA much sooner compared to D_2_Rs after an initial stimulant exposure may shift the balance in functional signaling and affect how reward circuits are activated with subsequent drug exposures *(18, 19)*. Furthermore, there is evidence that the enhancing and reinforcing effects of stimulant drugs may not only be mediated via D_2_Rs but also via D_1_Rs *(20)*. This difference in the neurochemical nature and timescale of D_1_ vs. D_2_R recycling has been unexplored in an *in vivo* setting as there have been no ready methods to measure these quantities in the living brain easily.

In the living brain, DA release can be measured non-invasively as a decrease in the *in vivo* binding of single-photon emission computed tomography (SPECT) and positron emission tomography (PET) imaging ligands such as [^123^I]IZBM, [^11^C]raclopride and [^11^C]PHNO *(8)*. PET studies of cocaine, amphetamine, and other stimulants have helped identify potential biomarkers that relate the concentration of D_2_R to compulsive patterns of drug use *(21, 22)*, and have shown that DA and striatal D_2_R are reduced in chronic drug abusers *(23, 24)*. Paradoxically, changes in receptor availability measured with PET following amphetamine stimulation persist well beyond acute fluctuations in extracellular DA concentrations, suggesting that mechanisms other than simple binding competition between DA and the PET ligand come into play *(25–28)*. Beyond PET imaging, discrepancies between microdialysis measurements of DA and hemodynamic responses have been attributed to receptor internalization *(29)*.

In this study, we investigate the internalization and recycling of D_1_R vs. D_2_R in nonhuman primates due to repeated amphetamine injections to depict discrepancies in intracellular mechanisms across DA receptor subtypes *in vivo*. We hypothesize that the D_1_R will be functionally active shortly after an amphetamine challenge, whereas the D_2_R will remain functionally inactive for up to 24h, as reported in previous *in vivo* PET studies. Combining amphetamine challenges with pharmacological blocking of D_1_Rs, functional responses of activated and subsequently internalized DA receptors were measured using simultaneous PET and functional magnetic resonance imaging (fMRI) in nonhuman primates.

## MATERIALS AND METHODS

### Study design

For the purpose of establishing the timeline of D_2_R internalization and brain-wide functional modulation due to repeated amphetamine, the experimental design consisted of two acute amphetamine administrations during two consecutive PET/MRI scans with the D_2_/D_3_ receptor PET radiotracer [^11^C]raclopride. In all studies, amphetamine (0.6 mg/kg bolus) was injected intravenously (i.v.) as a within-scan challenge approximately 40 min after starting a bolus-plus-infusion of the PET radiotracer, which enabled measuring dynamic signals across four states over time: The first scan provided a readout of the baseline state together with the effects of the first acute amphetamine injection (referred to as 0h), whereas the second scan evaluated the amphetamine-exposed state, as well as the effect of a second amphetamine injection either 3h or 24h later (Figure 1). We hypothesized that the excitatory D_1_R recycles faster and would therefore be available functionally at an earlier timepoint compared to the inhibitory D_2_R. We further hypothesized that amphetamine-induced excitatory D_1_-like and inhibitory D_2_-like receptor signaling manifest as positive and negative hemodynamic imaging signals, respectively. Therefore, to differentiate between D_1_ and D_2_R functional signaling, the D_1_R antagonist SCH 23390 (0.1 mg/kg + 0.09 mg/kg/h, bolus + infusion) was administered prior to the start of the PET/MR imaging session to block D_1_Rs in a subset of experiments.

**Figure 1.**
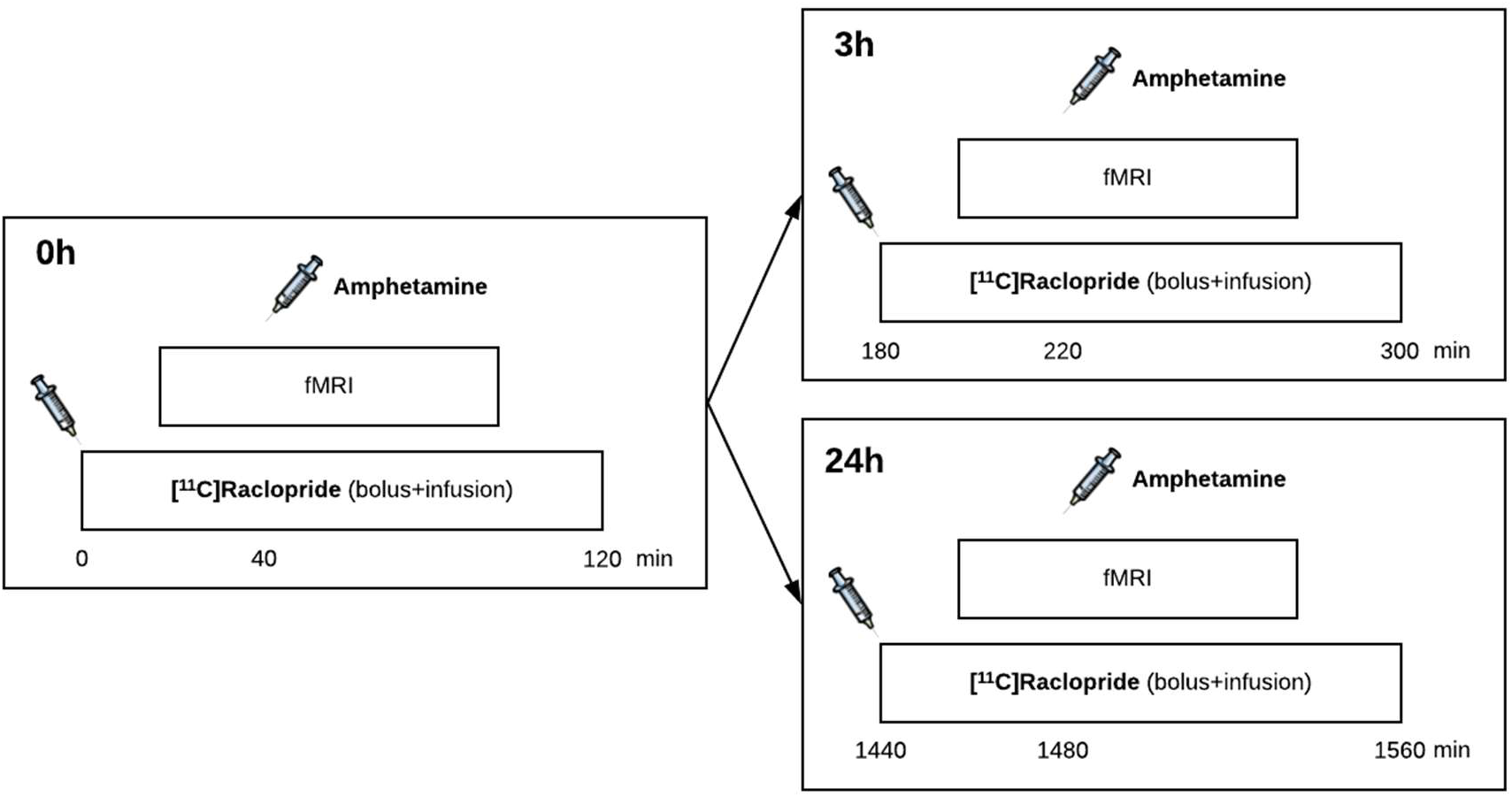
Schematic overview of the PET/MRI experiments. For each imaging session, the D_2_/D_3_ receptor PET radiotracer [^11^C]raclopride was administered using a bolus-plus-infusion paradigm, with fMRI acquired simultaneously throughout the scan. At 0h, an acute dose of amphetamine (0.6 mg/kg i.v. bolus) was administered 40 min after the injection of the radiotracer. This PET/MRI session was repeated in the same animal either 3h or 24h later, with a second amphetamine injection. In a subset of experiments, the D_1_ receptor antagonist SCH 23390 was administered as a bolus+infusion (0.1 mg/kg + 0.09 mg/kg/h) prior to the first [^11^C]raclopride injection.

### Animals

Three male rhesus macaques (8 (Animal 2), 14.5 (Animal 3), and 15 (Animal 1) years old) underwent PET/MRI. For each study, the animal was initially anesthetized with 10 mg/kg ketamine and 0.5 mg/kg xylazine, and then maintained with isoflurane (∼1%, mixed with oxygen) after intubation. Physiological parameters (blood pressure, pulse, end-tidal CO_2_, breathing rate, and oxygen saturation) were continuously monitored throughout the study. Animals were drug-free, i.e., had not undergone other pharmacological experiments at least one month prior to the experiments. All studies and procedures were approved by and complied with the regulations of the Institutional Animal Care and Use Committee at Massachusetts General Hospital.

### PET tracer injections

[^11^C]Raclopride was injected using a bolus+infusion protocol. Infusions employed *k*_bol_ values of 97.7 ± 26.6 min (n=9) for the [^11^C]raclopride injections at 0h, 83.1 ± 12.7 min (n=5) for the [^11^C]raclopride injections at 3h, and 95.7 ± 4.7 min (n=6) for the [^11^C]raclopride injections at 24 h. Boluses were administered by hand over a duration of 30 s, after which infusion at a rate of 0.01 ml/s was started with an automatic pump (Medrad Spectra Solaris). Specific activities at the time of injection were 1.49 ± 0.60 mCi/nmol (mean ± standard deviation), resulting in injected masses of 3.76 ± 1.66 μg on average. *Drugs:* Amphetamine (Sigma Aldrich, St Louis, MO, USA) was dissolved in saline immediately before the experiment and was administered as a slow bolus over 2 min. Amphetamine (0.6 mg/kg) was injected 39.3 ± 2.6 min (n = 15) after the injection of [^11^C]raclopride. SCH 23390 (Sigma Aldrich, St Louis, MO, USA) was administered using a bolus+infusion protocol to obtain a continuous blocking of D_1_R throughout the imaging session. SCH 23390 was administered 13.3 ± 3.6 min (n = 4) before the injection of [^11^C]raclopride using an MRidium infusion pump (IRadimed, Winter Springs, FL, USA). The bolus dose (0.1 mg/kg) was chosen based on previous NHP experiments*(61, 62)* and the infusion dose (0.09 mg/kg/h) was calculated based on a human subject [^11^C]SCH 23390 time-activity curve (TAC) and with the assumptions that metabolism is not changed from tracer dose to pharmacological dose and that SCH 23390 has similar kinetics in humans and nonhuman primates. For further information on the calculation of the *K*_bol_ for the SCH 23390 infusion, see Supplementary Materials.

### PET/MR Image Acquisition and Reconstruction

Simultaneous PET and MR data were acquired on a prototype scanner that consists of a BrainPET insert and a Tim Trio 3T MR scanner (Siemens AG, Healthcare Sector, Erlangen, Germany). A custom-built PET-compatible eight-channel NHP receive array *(63)* together with a vendor-supplied local circularly polarized transmit coil was used for MRI *(64)*. The phased array enabled two-fold acceleration with GRAPPA *(65)* in the anterior-posterior direction. Whole-brain fMRI data were acquired for the duration of the PET imaging using multi-slice echo-planar imaging (EPI) with an isotropic resolution of 1.3 mm and a temporal resolution of 3 s. Other parameters included FOV_MR_ = 110 × 72.8 mm^2^, BW = 1350 Hz per pixel, flip angle = 60° and an echo time of 23 ms. To improve fMRI detection power, ferumoxytol (Feraheme, AMAG Pharmaceuticals, Cambridge, MA) was injected at 10 mg/kg at the beginning of the fMRI acquisition *(66)*. No additional ferumoxytol was given for the imaging sessions at the 3h timepoint. In imaging sessions 24h later, the ferumoxytol dose was reduced to 8 mg/kg. PET emission data were acquired in list-mode format for 120 min (except for two scans where acquisition time was 100 min), starting with radiotracer injection. Images were reconstructed with a standard 3D Poisson ordered-subset expectation-maximization algorithm using prompt and variance-reduced random coincidence events. Normalization, scatter, and attenuation sinograms (including attenuation of the radiofrequency coil) were included in the reconstruction *(67)*. The reconstructed volumes consisted of 1.25 × 1.25 × 1.25 mm voxels in a 256 × 256 × 153 matrix, which were downsampled by a factor of two post-reconstruction. List-mode PET data were reconstructed into dynamic frames of increasing length (8 × 15 s, 8 × 30, 39 × 60, 10 × 120, 5 × 180, and 8 × 300 s).

### fMRI and PET data analysis

PET and MR data were registered to the Saleem-Logothetis stereotaxic space *(68)* with an affine transformation (12 degrees of freedom, DOF) using a multi-subject MRI template *(69)* in which standard regions of interest (ROI) were defined based on anatomy. To differentiate a more nuanced signal within the thalamus, we restricted the thalamus ROI to the thalamic region that encompassed the positive CBV signal at 0h. Furthermore, the paired 0h and 24h PET data were co-registered to obtain the best possible alignment of the ROIs.

Alignment of the EPI data used an affine transformation plus local distortion fields. After motion-correcting (AFNI software) and spatially smoothing fMRI data with a 2.5-mm Gaussian kernel, statistical analysis was carried out using the general linear model (GLM). Nuisance regressors corresponding to translations derived from the motion correction were included in the GLM analysis. The temporal response to the drug injection was modeled with a gamma variate function, in which the time to peak was adjusted to minimize the *χ*^2^/DOF of the GLM fit to the data. A long-lasting signal change that was distributed in several brain regions was modeled with a second gamma variate function. The resulting signal changes were converted to percent changes in CBV by standard methods *(70)*.

PET kinetic modeling employed a GLM formulation of the simplified reference tissue model (SRTM2) *(71)* with the cerebellum, excluding the vermis, as the reference region. For the quantification of binding changes over time due to the amphetamine interventions, the kinetic analysis included the time-dependent parameter *k*_2a_(t) *(72, 73)*, which was converted to a dynamic binding potential *(74)*. The reported pre-amphetamine BP_ND_s were calculated for the time-periods 0-40 min, 180-220 min, and 1440-1480 min. The reported post-amphetamine BP_ND_s were the dynamic BP_ND_ for the last time frame of the scan: 120 min, 300 min, and 1560 min.

All PET and fMRI data analysis and the generation of parametric images from voxelwise kinetic modeling were generated with open-access software (www.nitrc.org/projects/jip). Statistical values used for maps were computed by regularizing the random effects variance using an effective DOF of about 100 in the mixed-effects analysis *(75)*.

## RESULTS

### Amphetamine-induced receptor and functional maps across time

After each amphetamine injection, a reduction in [^11^C]raclopride-PET binding in the putamen and caudate was observed, driven by amphetamine-induced DA release. As seen in the upper row of Figure 2, the first amphetamine injection (at 0h) induced the largest decrease in D_2_R availability, while the second injections (at 3h and 24h) yielded smaller decreases compared to the 0h baseline. The reduction in D_2_R availability was quantified by changes in binding potential (ΔBP_ND_), from which D_2_R occupancy was determined (see paragraphs below). The upper row in the left panel of Figure 2 shows maps for changes in [^11^C]raclopride ΔBP_ND_ induced by each amphetamine injection alone. The right panel shows the equivalent maps for ΔBP_ND_ from experiments with a pre-block by the D_1_R antagonist SCH 23390. Corresponding whole-brain functional signaling was determined by simultaneous fMRI, and the parametric maps of negative (middle row) or positive (lower row) changes in cerebral blood volume (%CBV maps) from fMRI statistical analysis are shown in Figure 2. The use of an iron oxide contrast agent in this study enabled the conversion of fMRI signal changes to %CBV to quantify hemodynamic measures across sessions and represent drug-induced functional signaling. Amphetamine demonstrated both a positive and negative CBV component that was modulated and interestingly shifted in sign with repeated injections: The amphetamine challenge at 0h showed a predominantly negative CBV response localized to the putamen, caudate, thalamus, and cerebellum vermis (Figure 2). A small positive CBV signal was also observed bilaterally in the thalamus. The repeated amphetamine challenge 3h later resulted in a large positive CBV response localized to the putamen and caudate. The repeated amphetamine challenge 24h later elicited a response composed of both negative and positive responses localized in similar anatomical areas as described for the amphetamine challenge at 0h. However, the positive CBV response at 24h was much more pronounced in the striatum, similar to what was seen with the repeated amphetamine challenge after 3h.

**Figure 2.**
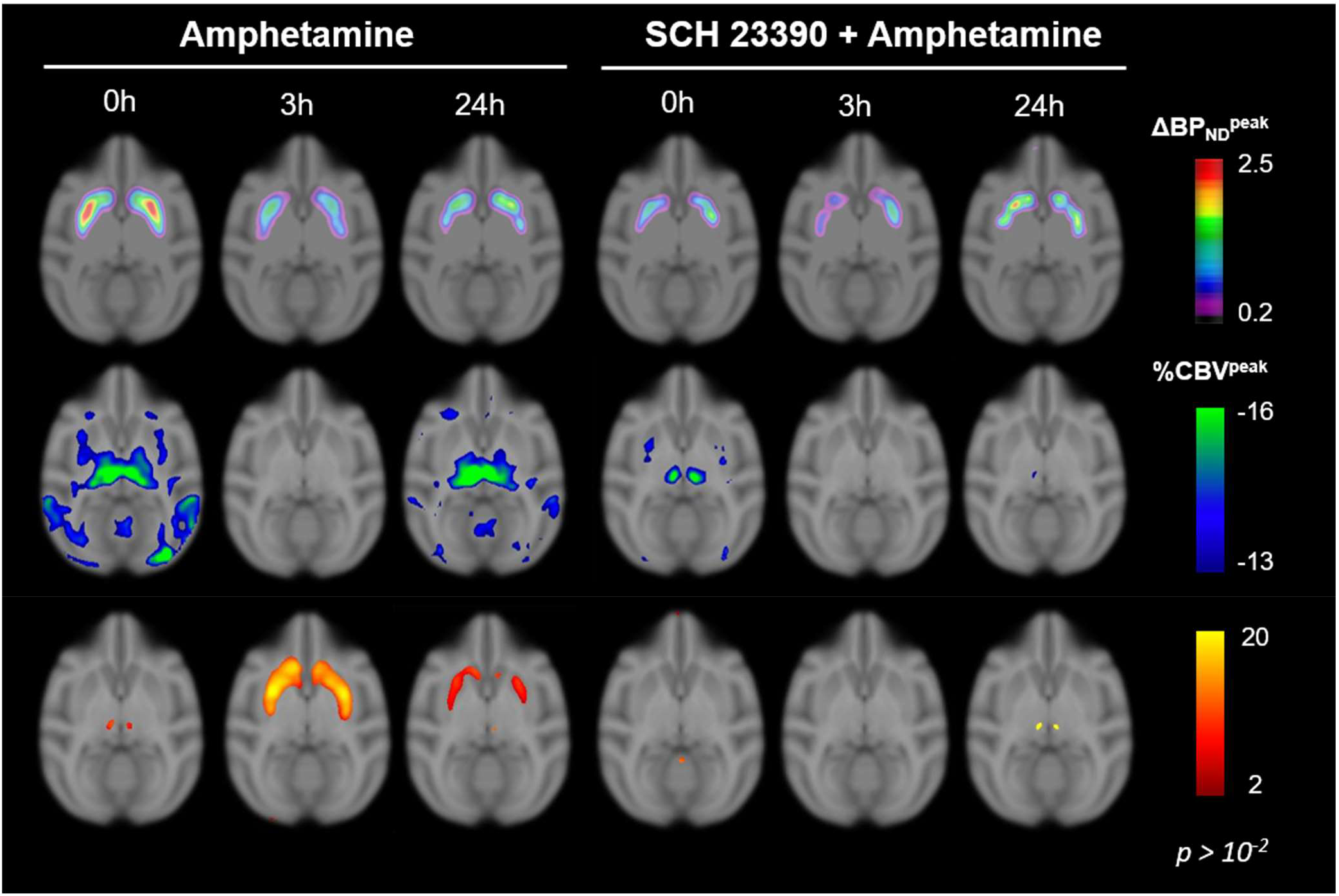
Parametric maps showing the change in [^11^C]raclopride binding potential (ΔBP_ND_) (upper row), together with simultaneously acquired percent changes in cerebral blood volume (%CBV) maps for the negative (middle row) and positive (lower row) peak response for the different experimental conditions: The first injection at 0h, followed by a second amphetamine injection either 3h or 24h, and the equivalent experiments with a pre-block by SCH 23390. Maps represent averages across repeated sessions in a total of three animals (see Methods for details). CBV maps were thresholded with a significance level of p < 10^−2^.

### Functional consequences of repeated amphetamine administration and D_2_R availability

Availability of long-lasting changes in baseline D_2_R availability was assessed with repeated scanning after 3h and 24h, and acute changes due to DA release were measured with each within-scan amphetamine administration. Figure 3 (upper row) shows average dynamic PET time-activity curves (TAC) for the putamen (a high-binding region) and cerebellum (the reference region) for the [^11^C]raclopride bolus+infusions at 0h, 3h, and 24h across all animals and sessions with amphetamine challenges (0.6 mg/kg, i.v.). TACs demonstrate an almost constant [^11^C]raclopride uptake around 30-40 min after radiotracer injection in both the high-binding and reference regions. Administration of amphetamine at 40 minutes resulted in the displacement of [^11^C]raclopride in the high binding regions putamen and caudate (Figure S1A-C in Supplementary Materials) at all three timepoints.

**Figure 3.**
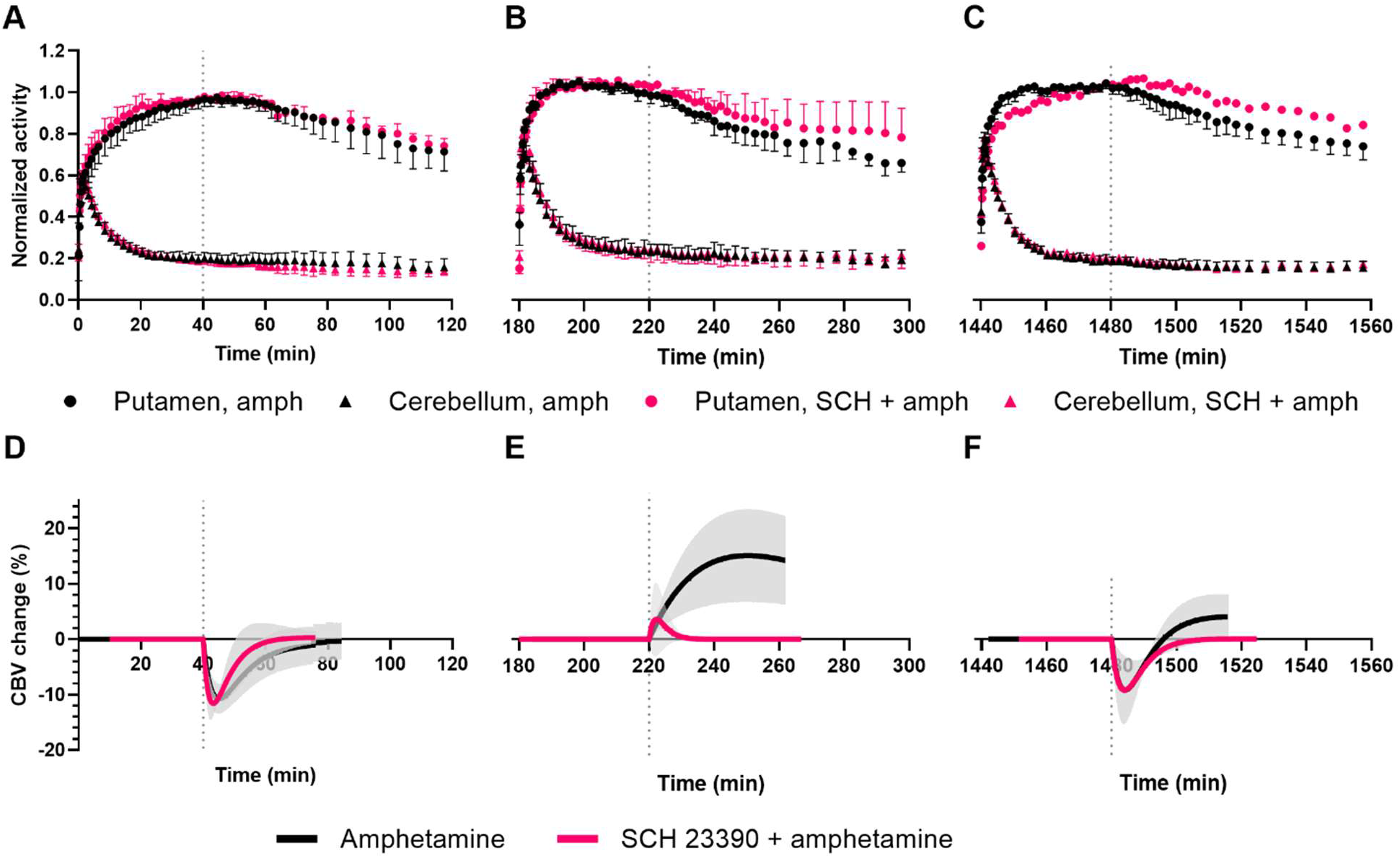
Mean time-activity curves for the putamen and cerebellum (normalized to peak Bq/mL value in the putamen) for the [^11^C]raclopride scans at 0h (A), 3h (B), and 24h (C). Mean timecourses for change in cerebral blood volume (CBV) in the putamen in response to the first amphetamine challenge (D), the second amphetamine challenge 3h later (E), and the amphetamine challenge 24h later (F). Black symbols represent experiments with amphetamine challenges (amph, 0.6 mg/kg), and pink symbols represent experiments in which the animals were pretreated with D_1_ receptor antagonist SCH 23390 (SCH, 0.1 mg/kg + 0.09 mg/kg/h) before the amphetamine (0.6 mg/kg) challenge. Vertical dotted lines represent the time of the amphetamine challenge at 40 minutes. Grey shaded areas represent standard deviation. Error bars represent standard deviation.

Despite only slightly reduced DA release observed from D_2_R occupancies at 3h, the CBV response was markedly different between the first (0h) and the repeated amphetamine challenge 3h later: The amphetamine-induced DA release at 0h caused a short-lasting decrease in CBV in the putamen (Figure 3D). The repeated amphetamine challenge 3h later caused a long-lasting increase in CBV in the putamen (Figure 3E). At 24h, the repeated amphetamine challenge caused a biphasic response with a negative CBV response similar to the 0h response and a positive longer-lasting component as seen with the amphetamine injection at 3h (Figure 3F).

Quantification of [^11^C]raclopride uptake in the high-binding regions before and after the drug challenges confirmed the amphetamine-induced decrease in binding (Figure 4A). The binding potentials (BP_ND_, mean ± SD) in the putamen decreased from 4.3 ± 0.8 to 3.2 ± 1.0 (n = 6) at 0h, from 3.2 ± 0.8 to 2.6 ± 0.6 (n = 3) at 3h and from 4.4 ± 0.8 to 3.8 ± 1.0 (n = 5) at 24h. As observed from the initial BP_ND_ before the amphetamine challenge in each session, D_2_R availability remained decreased for more than 3h, whereas it returned to baseline levels by 24h later. The corresponding peak D_2_R occupancies from the first amphetamine challenge (0h) was 27.3% [19.1; 35.1] (n = 6) (mean, [95% confidence interval]) in the putamen. The repeated amphetamine challenges 3h and 24h later resulted in slightly smaller occupancies of 19.3% [-9.4; 48.0] (n = 3) and 13.9% [2.2; 25.6] (n = 5) relative to the pre-amphetamine injection BP_ND_ in each session, respectively (Figure 4B and Figure S2). The lower occupancies at 3h and 24h suggest a reduced DA release capacity at these timepoints.

**Figure 4.**
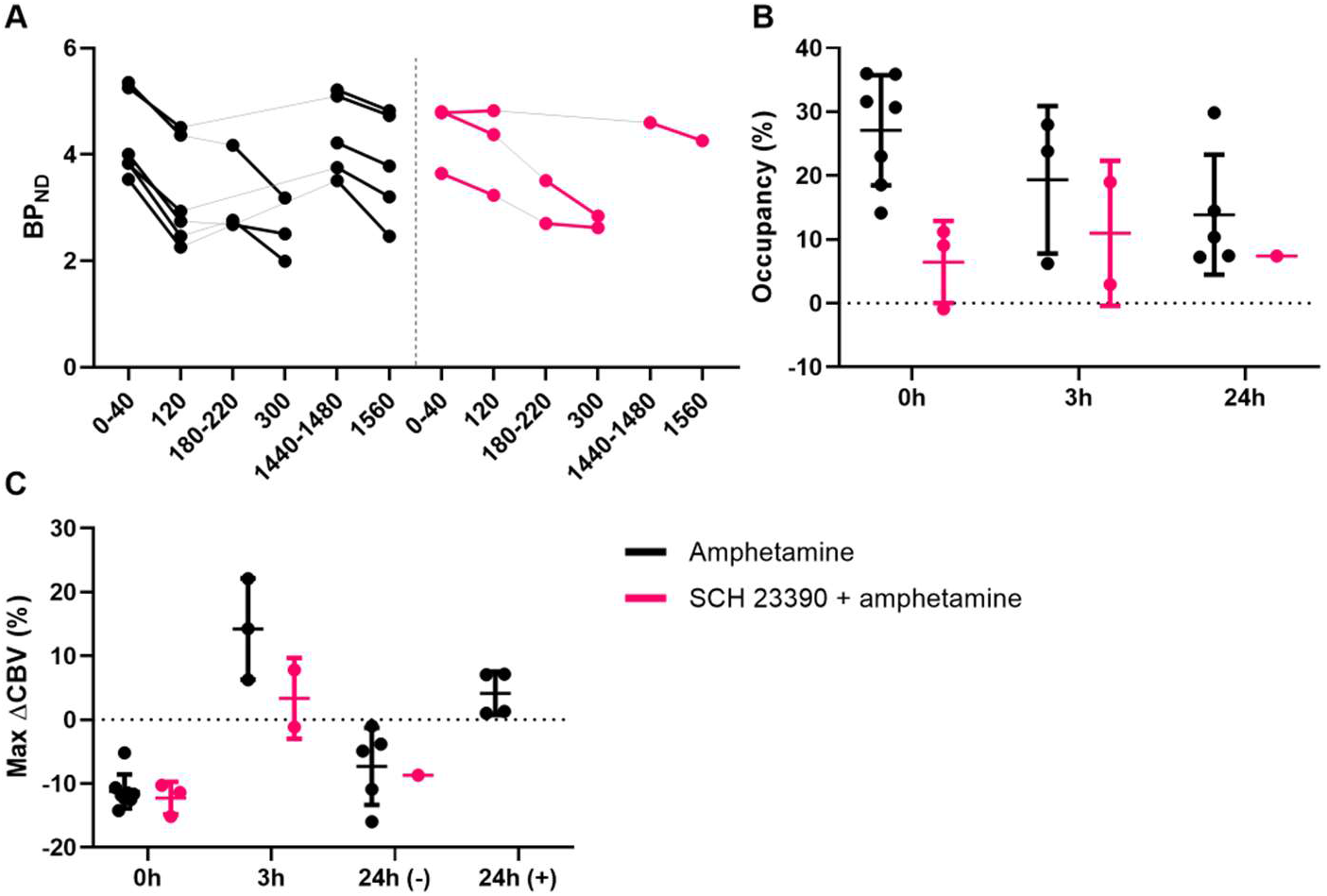
(A) Binding potentials (BP_ND_) in the putamen for all experiments before and after the amphetamine challenges. The x-axis denotes the time interval (in min) for each BP_ND_ calculation. (B) Peak occupancies due to each amphetamine challenge are calculated relative to their baseline within each session at the given timepoints. (C) Peak changes in CBV (ΔCBV) due to each amphetamine challenge. 24h (-) represents the peak changes in CBV of the negative response, whereas 24h (+) represents the peak changes in CBV of the positive response. Black symbols represent sessions with the administration of amphetamine only. Pink symbols represent sessions in which D_1_Rs were blocked by SCH 23390 before the start of the PET/MR acquisition. Error bars represent standard deviation.

The peak CBV responses from the general linear model fit to the measured data are shown in Figure 4C. The peak %CBV changes [95% confidence interval] of the amphetamine challenge at 0h was -11.2 [-13.4; -9.0] (n = 8), whereas the peak %CBV changes of the repeated amphetamine challenge 3h later was 14.2 [-5.4; 33.9] (n = 3). The peak %CBV changes of the repeated amphetamine challenge 24h later was biphasic with a short-lasting negative component (−7.3 [-14.8; 0.2], n = 5) and a longer-lasting positive component (4.1 [-1.3; 9.6], n = 5).

Combined, these PET and fMRI timecourses show that DA release occurred with each amphetamine challenge in the striatum, however, with slightly reduced DA release capacity at 3h and 24h. D_2_R availability remained reduced at 3h but returned to baseline levels after 24h. Most strikingly, the CBV response in the striatum inverted from a short-lasting, predominantly inhibitory response at 0h to a long-lasting excitatory response at 3h. After 24h, the CBV response returned to a short-lasting negative response.

### Amphetamine responses during D_1_R blockade

We hypothesized that the positive CBV response induced by the repeated amphetamine challenge 3h later was a consequence of activating excitatory D_1_Rs. To test this hypothesis, we blocked D_1_Rs with the antagonist SCH 23390.

Although TACs appeared strikingly similar with or without SCH 23390 pretreatment (Figure 3A-C, pink series), quantification of radiotracer pharmacokinetics before and after the challenges showed smaller amphetamine-induced decreases in binding (Figure 4A). The binding potentials (BP_ND_, mean ± SD) in the putamen decreased from 4.4 ± 0.7 to 4.1 ± 0.8 (n = 3) at 0h, from 3.1 ± 0.6 to 2.7 ± 0.2 (n = 2) at 3h and from 4.6 to 4.0 (n = 1) at 24h. Thus, the kinetic modelling revealed a lower peak occupancy in the putamen of -0.9%, 9.1%, and 11.2% at 0h (n = 3), 19.0% and 2.9% at 3h (n = 2) and 7.4% at 24h (n = 1) with SCH 23390 as a D_1_R blocker (Figure S2A-C in Supplementary Materials).

Under D_1_R blocking, the timecourse of the CBV response induced by the 0h amphetamine challenge was a predominantly negative response, similar to the non-blocked condition (Figure 3D) and in line with inhibitory signaling. However, the repeated amphetamine challenge 3h later induced only a small and short-lasting increase in CBV (Figure 3E), which was very different from the large positive CBV response seen in the non-blocked condition. The CBV signal at 24h resembled the 0h amphetamine response with a negative CBV and no positive component (Figure 3F). The CBV responses were quantified (Figure 4C), with peak %CBV changes [95% confidence intervals] after pretreatment with SCH 23390 being -12.3 [-18.6; -5.9] at 0h (n = 3), 7.8 and -1.1 at 3h (n = 2) and -8.7 at 24h (n = 1) in the putamen.

The ΔBP_ND_ and CBV maps confirm that voxelwise CBV responses were reduced in magnitude after SCH 23390 pretreatment. The positive CBV response at the repeated amphetamine challenges 3h and 24h later was abolished entirely (Figure 2).

### Regional differences in amphetamine-induced signaling

The CBV timecourses in the caudate and nucleus accumbens in response to the amphetamine challenges at 0h, 3h, and 24h with and without SCH 23390 pretreatment (Figure 5) were similar to the responses observed in the putamen (Figure 3D-F), both in shape and magnitude. The first amphetamine injection at 0h yielded a predominantly negative signal in all regions, except in the thalamus, where a prominent biphasic CBV response was observed at both 0h and 24h. While the CBV response inverted to a dominant positive response in the striatal regions at 3h, the thalamus exhibited a more moderate positive CBV signal than the striatal regions. Interestingly, the positive thalamic CBV response was eliminated at 0h and 3h by SCH 23390 pretreatment, but not at 24h – contrary to the positive striatal component, which was fully blocked by SCH 23390 at all timepoints.

**Figure 5.**
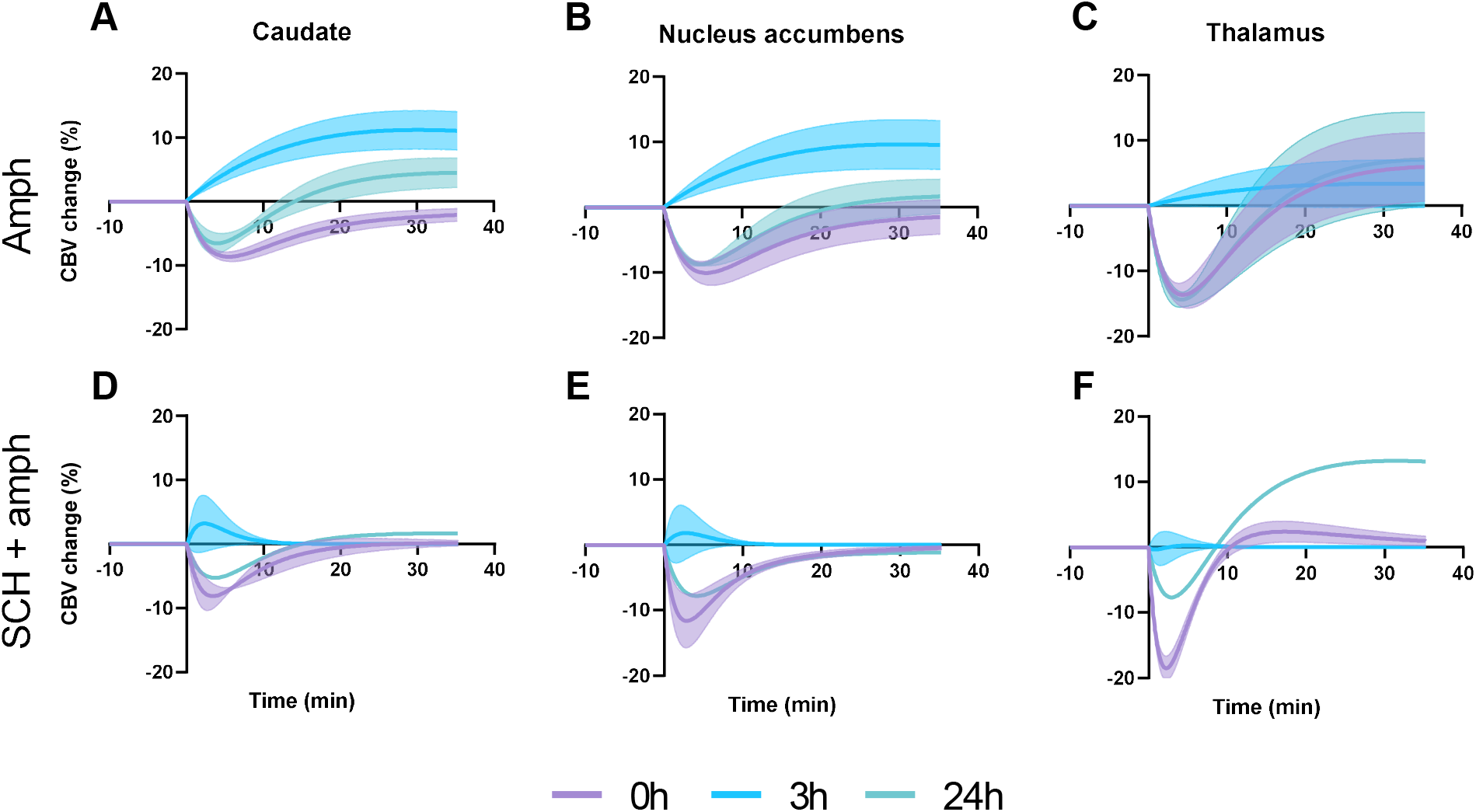
Cerebral blood volume (CBV) timecourses in percent change from baseline, shown as the mean of the GLM fit, in response to amphetamine (upper row) and after pretreatment with SCH 233390 (lower row) in the caudate (A, D), nucleus accumbens (B, E) and thalamus (C, F). The first amphetamine injection at 0h is overlayed with the start of the second amphetamine injections at 3h or 24h for comparison purposes. Shaded areas represent standard errors of the mean.

## DISCUSSION

This study shows a temporal discrepancy in D_1_ vs. D_2_/D_3_ receptor recycling in the living brain in nonhuman primates. Although both receptor subtype classes may internalize in response to an amphetamine challenge, our results indicate that the D_1_R subtype quickly recycles and is available for reactivation 3h and 24h after an acute amphetamine challenge. In contrast, D_2_/D_3_ receptors stay internalized and non-functional for more than 3h with a return to functionality by 24h.

A single amphetamine challenge induced endogenous DA release, which decreased [^11^C]raclopride binding together with CBV, as measured with simultaneous PET/fMRI. After 3h, D_2_/D_3_ receptor availability remained reduced beyond the expected timeline of DA release *(30)*, consistent with D_2_/D_3_ receptors being internalized. Supporting this further, we found that the initial negative CBV response, driven by activation of inhibitory D_2_/D_3_ receptors after a single amphetamine injection, was not predominant at 3h after a second amphetamine injection. The lack of a negative CBV component suggests that the D_2_/D_3_ receptors reported by [^11^C]raclopride-PET are non-functional at this early time point after prior exposure. Despite the lower availability of D_2_/D_3_ receptors 3h later, we found that DA release capacity was unchanged. With a proportion of the D_2_/D_3_ receptors internalized and non-functional, the repeated amphetamine-induced DA surge instead resulted in activating excitatory D_1_Rs and thereby increasing CBV. This activation was confirmed to be D_1_R-mediated by blocking the D_1_Rs with the antagonist SCH 23390, abolishing the positive CBV response altogether. This modulation of fMRI signal over time is coherent with a shift from D_2_R-driven inhibitory signaling at 0h to D_1_R-driven excitatory signaling at 3h.

When the amphetamine challenge was repeated 24h later, DA release capacity was comparable to the amphetamine challenge at 0h. The negative CBV response that was present at the 0h amphetamine injection had been reestablished, suggesting that the D_2_/D_3_ receptors had recycled to the cell membrane surface and were yet again functional. The positive CBV component that dominated the response at 3h persisted at 24h and could be blocked by SCH 23390 in the DA-rich striatum. Contrary to that, the (positive part of the) biphasic signal in the thalamus could not be blocked by SCH 23390 at 24h, suggesting that the excitatory thalamic component was not DA-mediated. It points to striatal-thalamic signaling, which was modulated differentially at 24h. Specifically, this type of neuroanatomical interaction exists between local D_1_R-mediated transmission in the striatum and excitatory glutamatergic projections from the thalamus *(31)*. It appears to be critical in relapse to methamphetamine seeking after prolonged withdrawal.

Several studies report that D_2_R internalization is dependent on β-arrestin2 (also referred to as arrestin3) *(12, 32, 33)*. Furthermore, D_2_Rs are targeted by lysosomes for degradation via interaction with G protein-coupled receptor (GPCR) associated sorting protein *(16)*. Skinbjerg et al. showed that an amphetamine challenge reduced binding potentials 4h after an amphetamine challenge in wild-type mice but not in β-arrestin knock-out mice, indicating that internalization and recycling are β-arrestin-dependent*(27)*. The study also demonstrates that internalization is the driving mechanism for the temporal discrepancy between the DA microdialysis measures and the long-lasting decrease in radiotracer binding (as described in the introduction). Other studies have also shown that [^123^I]IBZM and [^11^C]raclopride binding potentials remain reduced 3h and 24h after a single amphetamine challenge *(25, 26)*. While the PET-based results in the latter studies concur with our findings, the discrepancy between receptor availability and functionality we report indicates that PET imaging alone may not always fully capture the state of receptors after agonist exposure.

The D_1_Rs undergo classical GPCR regulation, rapidly desensitizing and internalizing via GPCR kinase phosphorylation *(34)*, β-arrestin 2 binding, and clathrin-mediated endocytosis *(35)*. Following endocytosis, the D_1_Rs are resensitized and recycled back to the plasma membrane where they can bind ligand once again*(36–38)*. An *in vivo* study showed that D_1_Rs internalized rapidly but remained in intracellular compartments for more than 90 min following an amphetamine challenge *(14)*. In cell cultures, D_1_R mediated cAMP production, i.e., resensitization, returned to baseline 5-6h after agonist exposure *(37)*. In NHPs, a 5-7% decrease in BP_ND_s was found 2h after a high dose amphetamine (2.0 mg/kg) challenge measured by two different D_1_R selective PET radiotracers*(39)*, suggesting that recycling of the receptors could occur relatively quickly. While not all PET radiotracers are sensitive to changes in neurotransmitter levels, the latter study benefits from having investigated the amphetamine-induced D_1_R recycling with both an antagonist and agonist radiotracer. Given these results, it seems likely that the D_1_Rs are available for functional activation 3h after the amphetamine challenge, as seen in the present study. Bartlett et al. found a discrepancy in the cellular recycling of D_1_R and D_2_Rs, where D_1_Rs were found to recycle to the plasma membrane. In contrast, D_2_Rs were targeted for degradation *(16)*, supporting a temporal discrepancy in D_1_R and D_2_/D_3_ receptor recycling. These data aligned well with the fMRI data measured in our study: the functional response changed from being driven by D_2_Rs to being dominated by D_1_Rs after 3h because the D_2_Rs were internalized after the initial amphetamine challenge.

Our study confirms that [^11^C]raclopride binding potentials returned to baseline by 24h after amphetamine exposure *(26)*. We found a slight decrease in amphetamine-induced D_2_/D_3_ receptor occupancy at 24h, i.e., reduced DA release, which could result from decreased DA synthesis. Reduced DA synthesis capacity in cocaine users as measured with [^18^F]DOPA support this hypothesis *(40)*. The CBV response at the repeated amphetamine challenge 24h later was biphasic and resembled a mix of CBV responses of the 0h and the repeated amphetamine challenge 3h later. This can be interpreted as the resumption of D_2_/D_3_ receptor functionality, either by de novo receptor synthesis or recycling of receptors from intracellular compartments.

Because DA has a higher affinity for the D_2_R than for the D_1_R, the amount of DA release can drive the balance between excitatory D_1_R and inhibitory D_2_R signaling, with the combination of both making up the fMRI signal. A model for DA-induced fMRI signal has previously described how the *in vivo* functional response to DA results in a biphasic response with an initial D_2_R-driven negative followed by a D_1_R-driven positive component *(41)*. Our CBV signals from the initial amphetamine challenge match this model, and we further demonstrate experimental *in vivo* evidence of how the balance between D_1_ and D_2_R signaling can affect responses to repeated amphetamine. The initial short-lasting decrease in CBV also mimics the response induced by the D_2_R selective agonist quinpirole, which we have previously shown to be a signature of rapid D_2_R desensitization and internalization *(42)*. In concordance with such a model, the fMRI temporal profile at 3h with the repeated amphetamine challenge in this study is remarkably similar to a predominantly D_1_R activation, i.e., with D_2_/D_3_ receptors internalized. A complementary interpretation to the varying recovery times is that D_1_Rs may not be internalized to the same extent due to the lower affinity of DA for D_1_R relative to D_2_R.

Microdialysis and fast-scan cyclic voltammetry studies have shown that synaptic DA levels return to baseline 2-3 hours after an amphetamine challenge *(43–45)*. This is in line with our findings, where the 0h and repeated amphetamine challenge 3h later resulted in comparable D_2_/D_3_ receptor occupancies, suggesting that vesicle DA concentration had been restored and that the DA release capacity was unchanged. Only in the sessions with SCH 23390 pretreatment, we observed lower amphetamine-induced D_2_R occupancies. Serotonin 5-HT_2A_ receptor antagonism has been shown to attenuate amphetamine-elicited DA release without affecting basal DA levels *(46, 47)*. This is relevant because SCH 23390 also binds, albeit with lesser affinity, to 5-HT_2A_ receptors and thus may explain the reduced DA release during SCH 23390 blockage. Another reason may be the ability of SCH 23390 to increase extracellular DA levels *(48, 49)* and consequently decrease DA release capacity. A small DA release induced by SCH 23390 may also explain why baseline binding potentials were lower at the 0h amphetamine challenge with SCH 23390 pretreatment compared to the non-pretreated session.

Imaging genetically modified animals that cannot internalize D_2_/D_3_ receptors and comparing their CBV-occupancy timecourses to wild-type animals would provide more direct evidence of receptor internalization. Alternatively, treatment with β-arrestin inhibitors such as barbadin *(50)* can offer a pharmacological approach to deciphering internalization. Investigating receptor internalization mechanisms in animal models of substance abuse would be highly relevant and could help identify biomarkers to guide treatment.

D_1_R and D_2_Rs have been shown to mediate opposing effects on drug-seeking behavior. While stimulation of D_2_R induces stimulant-seeking behavior, stimulation of D_1_Rs attenuates it, possibly by satiating reward pathways *(51)*. Importantly, since blocking either D_1_R or D_2_Rs has been shown to attenuate reinstatement of cocaine-seeking, both receptors seem to play a crucial role in drug-seeking responses [reviewed by Self et al. *(18)*]. Our results delineate this intricate interplay between D_1_R and D_2_R and point to differential receptor externalization times as a mechanism affecting the functional signaling to repeated doses of amphetamine. Systemic administration of the D_1_R antagonist SCH 23390 reduces multiple addiction-related behaviors, including reward, self-administration, and priming-induced drug seeking *(52–54)*. A recent study showed that methamphetamine self-administration enhances the expression of D_1_R internalization-promoting proteins in the dorsal striatum, whereas SCH 23390 reduces this effect *(55, 56)*. In this way, stimulants may alter D_1_R responsiveness to DA surges and regulate the reinforcing effects of the drugs. Reduced long-term potentiation after methamphetamine administration has also been measured *(55)*, which may manifest as reduced CBV in the dorsal striatum as observed here. Together, these findings suggest that pharmacological blocking of D_1_Rs in the dorsal striatum reduces stimulant intake and rescues stimulant-induced depression of synaptic plasticity.

Clinically, amphetamine and other drugs that act on DA transporters are used in the treatment of attention deficit hyperactivity disorder and narcolepsy. Both D_1_Rs and D_2_Rs likely mediate the pro-attentional effects of DA-elevating drugs. However, recent evidence points towards a crucial role of the D_1_Rs. Administration of a D_1_R partial agonist improved attention/vigilance in rats during a demanding task *(57)*. Similarly, amphetamine also enhanced performance in a 5-choice continuous performance test in humans, rats, and mice *(58, 59)*. Importantly, this effect was observed irrespective of acute treatment with the D_2_R antagonist haloperidol in rats, suggesting that the pro-attentional effects of amphetamine are predominantly a D_1_R-mediated mechanism. This appears compatible with patients’ concurrent use of antipsychotics since these drugs have a lower affinity for the D_1_R than the D_2_R *(60)*. Preclinical and clinical data show that low striatal D_2_R availability is associated with increased drug self-administration and impulsive behavioral patterns *(2)*. Although existing data support the view that impulsivity is a predictive phenotype for addiction, it is still unknown whether the reduced DA transmission is a consequence of drug abuse or an underlying vulnerability factor for substance abuse. Given the present study results, we speculate that patients with substance use disorder have augmented D_2_R internalization time and consequently would be at higher risk of impulsive behaviors leading to a preference for small immediate rewards and increased drug self-administration. Nevertheless, it would be highly relevant to investigate a patient population with a high risk for developing substance use disorder using a similar experimental design.

In conclusion, amphetamine-induced DA release activates all DA receptors upon which they internalize. Our data provide *in vivo* evidence for a temporal discrepancy between D_1_ and D_2_/D_3_ receptor recycling. This finding had previously only been indicated *in vitro* in the rodent brain. The present study extends these findings into the primate brain *in vivo* in the context of repeated amphetamine challenges. Inhibitory D_2_/D_3_ receptors drive the functional response to an initial amphetamine challenge, whereas the functional response to a repeated amphetamine challenge 3h later is dominated by excitatory D_1_Rs. D_1_Rs are likely not internalized to the same degree or recycle to the cell membrane surface faster than the D_2_/D_3_ receptors. Pharmacological blocking of the D_1_Rs or preventing the internalization and degradation of D_2_Rs could restore the balance between D_1_R vs. D_2_R signaling. This may be a potential therapeutic avenue in treating substance use disorders. Overall, these results contribute to the mechanistic understanding of how stimulants modulate the dopaminergic system and how this may ultimately lead to substance use disorder.

## Supplementary information is available

Table S1

Figure S1-S2

Supplementary Methods: Calculating the bolus + infusion ratio of SCH 23390

## Acknowledgments

The authors would like to thank the Radiochemistry and integrated PET/MR team at the Athinoula A. Martinos Center for Biomedical Imaging, Department of Radiology, Massachusetts General Hospital. The authors would also like to thank Dr. Joseph Coyle for helpful comments on the manuscript and Dr. Brice Ozenne (Department of Public Health, Section of Biostatistics, University of Copenhagen and Neurobiology Research Unit, Copenhagen University Hospital, Copenhagen, Denmark) for reading the manuscript and providing statistical consulting.

## Author contributions

Conceptualization: HDH, CYS

Methodology: HDH, MS, CYS

Software: JBM, CYS

Formal analysis: HDH, CYS

Investigation: HDH, HPD, CYS

Writing – original draft: HDH, CYS

Writing – review & editing: HDH, MS, HPD, JBM, BRR, CYS

Funding acquisition: CYS

## Data materials availability

Imaging data and code used in the analysis are available upon request.

