## Supplementary material for "Differential D_1_ and D_2_ receptor internalization and recycling induced by amphetamine *in vivo*"

**Table S1:** Overview of the number of scanning sessions at the 0h, 3h, and 24h timepoints with amphetamine injection or SCH 23390 blockage in combination with amphetamine. A total of three non-human primates were scanned across the experimental conditions. The sessions correspond to Figure 2 of the manuscript.

|  | **Amphetamine** | | | **SCH 23390 + amphetamine** | | |
| --- | --- | --- | --- | --- | --- | --- |
|  | **0h** | **3h** | **24h** | **0h** | **3h** | **24h** |
| **PET** | n=6 | n=3 | n=5 | n=3 | n=2 | n=1 |
| **fMRI** | n=8 | n=3 | n=5 | n=3 | n=2 | n=1 |

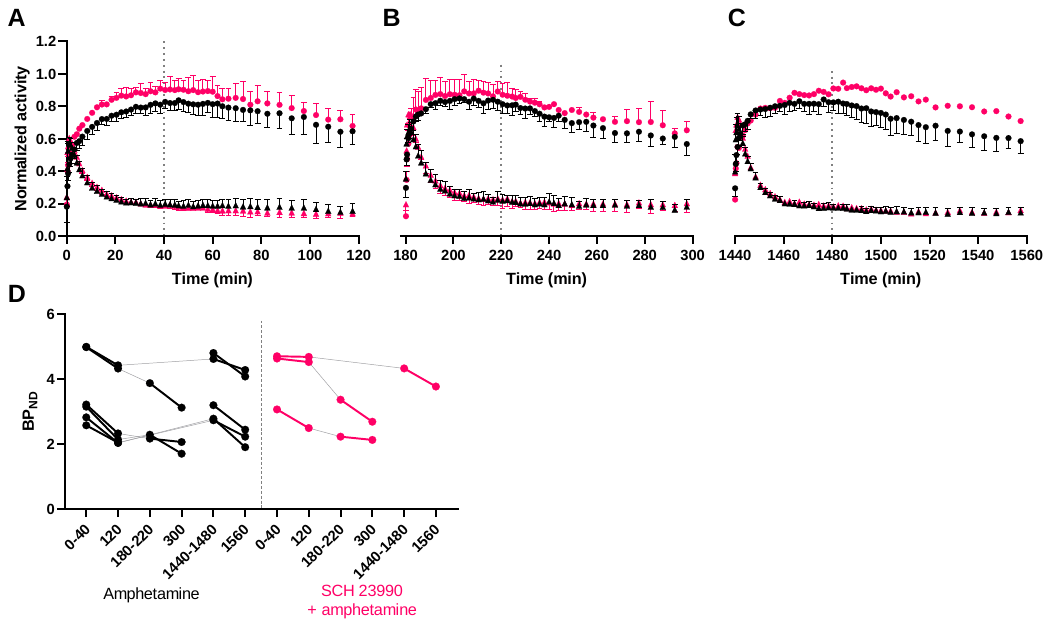
 **Figure S1.** Mean time-activity curves for the caudate and cerebellum (normalized to peak Bq/mL value in the putamen) for the [^11^C]raclopride scans at 0h (A), 3h (B), and 24h (C). Binding potentials (BP_ND_) in the caudate for all experiment before and after the amphetamine challenges (D). Numbers on the x-axis denotes the time-interval for which each BP_ND_ was calculated. Black symbols represent experiments with amphetamine (amph, 0.6 mg/kg) challenges whereas pink symbols represent experiments in which the animals were pretreated with D_1_ receptor antagonist SCH 23390 (SCH, 0.1 mg/kg + 0.09 mg/kg/h) before the amphetamine (0.6 mg/kg) challenge. Vertical dotted lines represent time of amphetamine challenge. Grey lines connect paired experiments. Error bars represent standard deviation.

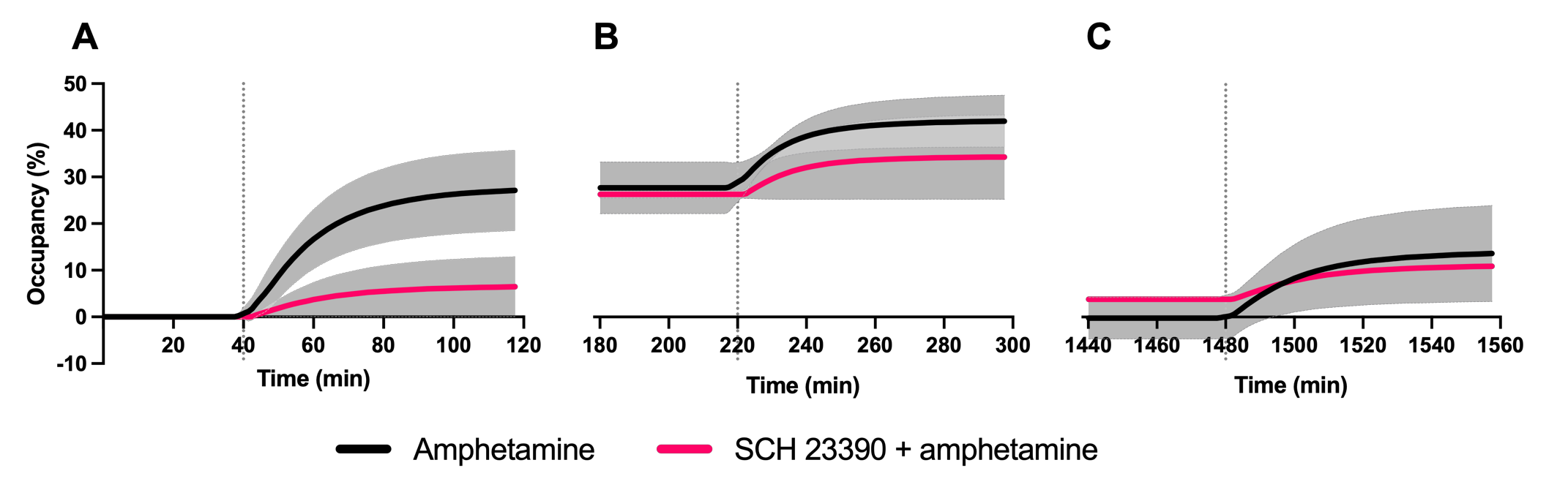

**Figure S2.** Mean occupancy timecourses in the putamen in response to the (**A**) first amphetamine challenge, (**B**) the second amphetamine challenge 3h later, or (**C**) the amphetamine challenge 24h later. Black symbols represent experiments with administration of 0.6 mg/kg amphetamine. Pink symbols represent a session in which SCH 23390 (0.1 mg/kg + 0.09 mg/kg/h) was administered before the start of the PET-MR acquisition. Error bars represent standard deviation.

**Supplementary Methods: Calculating the bolus + infusion ratio of SCH 23390**

Based on a human subject time-activity curve (TAC) and a predicted bolus length of 5 sec, a set of input functions was calculated to determine which one would best predict steady-state levels of SCH23390 in the striatum. Different infusion fractions α were tested and it was found that an infusion fraction of α = 0.005 to 0.006 would achieve steady-state levels in the striatum. We then determined the magnitude of the bolus component for the bolus + infusion rate (*K*_bol_, which defines “how many minutes worth of infusion the bolus corresponds to” or “for how long we need to infuse for the infusion to match the bolus”) as *K*_bol_ = 𝑇_𝑏𝑜𝑙𝑢𝑠_ α. Given a bolus injection time of 𝑇_𝑏𝑜𝑙𝑢𝑠_ = 20 seconds and an infusion fraction of α = 0.005, this resulted in a *K*_bol_ = 67 minutes.
